# Transcriptional survey of ovarian bacteriomes in the cereal weevil, *Sitophilus oryzae*, shows down-regulation of immune effectors at the onset of sexual maturity

**DOI:** 10.1101/2023.01.31.526389

**Authors:** Mariana Galvão Ferrarini, Agnès Vallier, Elisa Dell’Aglio, Séverine Balmand, Carole Vincent-Monégat, Mériem Debbache, Justin Maire, Nicolas Parisot, Anna Zaidman-Rémy, Abdelaziz Heddi, Rita Rebollo

**Affiliations:** Univ Lyon, INRAE, INSA-Lyon, BF2I, UMR 203, 69621 Villeurbanne, France; Université de Lyon, Université Lyon 1, CNRS, Laboratoire de Biométrie et Biologie Evolutive UMR 5558, F-69622 Villeurbanne, France; Univ Lyon, INSA-Lyon, INRAE, BF2I, UMR 203, 69621 Villeurbanne, France

**Keywords:** Coleoptera, symbiosis, bacteria, bacteriome, interaction

## Abstract

Insects often establish long-term relationships with intracellular symbiotic bacteria, *i.e*. endosymbionts, that provide them with essential nutrients such as amino acids and vitamins. Endosymbionts are typically confined within specialized host cells called bacteriocytes that may form an organ, the bacteriome. Compartmentalization within host cells is paramount for protecting the endosymbionts and also avoiding chronic activation of the host immune system. In the cereal weevil *Sitophilus oryzae*, bacteriomes are present as a single organ at the larval foregut-midgut junction, and in adults, at the apex of midgut mesenteric caeca and at the apex of the four ovarioles. While the adult midgut endosymbionts experience a drastic proliferation during early adulthood followed by complete elimination through apoptosis and autophagy, ovarian endosymbionts are maintained throughout the weevil lifetime by unknown mechanisms. Bacteria present in ovarian bacteriomes are thought to be involved in the maternal transmission of endosymbionts through infection of the female germline, but the exact mode of transmission is not fully understood. Here, we show that endosymbionts are able to colonize the germarium in one-week-old females, pinpointing a potential infection route of oocytes. To identify potential immune regulators of ovarian endosymbionts, we have analyzed the transcriptomes of the ovarian bacteriomes through young adult development, from one-day-old adults to sexually mature ones. In contrast with midgut bacteriomes, immune effectors are downregulated in ovarian bacteriomes at the onset of sexual maturation. We hypothesize that relaxation of endosymbiont control by antimicrobial peptides might allow bacterial migration and potential oocyte infection, ensuring endosymbiont transmission.

## Introduction

Symbiosis is a widespread phenomenon in nature. Insects dwelling in unbalanced diets have recurrently established long-term associations with intracellular symbiotic bacteria (endosymbionts) that provide them with essential nutrients, such as amino acids or vitamins (Moran et al., 2008; Moya et al., 2008). Such associations allow insects to invade and colonize new environments otherwise inhabitable, increasing their fitness and hence their socioeconomic burden. For instance, bloodsucking insects, including *Cimex lectularius* (bedbugs) and *Glossina spp*. (Tsetse fly), rely on B vitamins provided by their endosymbionts, *Wolbachia* (Hosokawa et al., 2010) and *Wigglesworthia glossinidia* (Snyder et al., 2010) respectively, while stored-product pest *Sitophilus spp*. (cereal weevils) relies on the Gram-negative bacterium *Sodalis pierantonius* which provides aromatic amino acids as precursors for insect cuticle synthesis (Heddi et al., 1998; Oakeson et al., 2014; Vigneron et al., 2014).

Endosymbionts are generally contained within specialized host cells, the bacteriocytes, that may form an organ, the bacteriome (Pierantoni, 1927; Buchner and Mueller, 1965; Nardon et al., 2002). Confinement of endosymbionts within bacteriocytes ensures their survival in a particular immune micro-environment and avoids chronic activation of the host immune system (Masson et al., 2015b; Maire et al., 2019; Ferrarini et al., 2022). This is particularly important in recently established relationships where endosymbionts encode virulence-related genes, such as in the *Sitophilus* spp./*S. pierantonius* association, where the endosymbiont’s genome encodes and expresses genes involved in both type III secretion system and in Microbial-Associated Molecular Patterns (MAMPs), such as peptidoglycans (Dale et al., 2002; Oakeson et al., 2014; Maire et al., 2019, 2020b). Artificial hemolymph infection of *S. pierantonius* results in a systemic immune activation associated with the production of a cocktail of host antimicrobial peptides (AMP) (Anselme et al., 2008; Ferrarini et al., 2022). Within bacteriomes, *S. pierantonius* is permanently targeted by the AMP Coleoptericin A (ColA), which interacts with the chaperonin GroEL, and hence inhibits bacterial cell division (Login et al., 2011). Expression of the bacteriome-specific AMP ColA at standard conditions relies on the transcription factor *relish*, and the immune-deficiency (IMD) pathway but not on the peptidoglycan recognition protein LC receptor (PGRP-LC) (Maire et al., 2018, 2019). In addition to ColA, PGRP-LB is also expressed within weevil’s bacteriomes and acts as the ultimate safeguard, by cleaving a monomeric form of peptidoglycan (tracheal cytotoxin, TCT) therefore preventing leakage from the bacteriome and subsequent chronic activation of an IMD-dependent immune response (Maire et al., 2019). Altogether, these molecular mechanisms provide bacteriomes with the capacity to protect endosymbiotic bacteria and maintain symbiotic benefits, while preserving the host homeostasis.

In the cereal weevil *Sitophilus oryzae*, endosymbionts are housed within a bacteriome located at the larval foregut-midgut junction, in many bacteriomes at the apex of adult midgut caeca, as well as at the apex of female ovaries (Figure 1A). In young adult weevils, an exponential increase in endosymbionts was observed in midgut bacteriomes during the first week of adulthood (Vigneron et al., 2014), triggered by the host carbohydrate intake (Dell’Aglio et al., 2022). The drastic increase in endosymbiont load is accompanied by an increase in the bacteriome-specific AMP ColA (Masson et al., 2015a), and was shown to provide aromatic amino acids necessary for the host cuticle maturation (Vigneron et al., 2014). Following two weeks of adult life, endosymbionts are eliminated from the midgut bacteriomes through apoptosis and autophagy, while ovarian endosymbionts are kept through the weevil’s lifetime (Vigneron et al., 2014).

**Figure 1.**
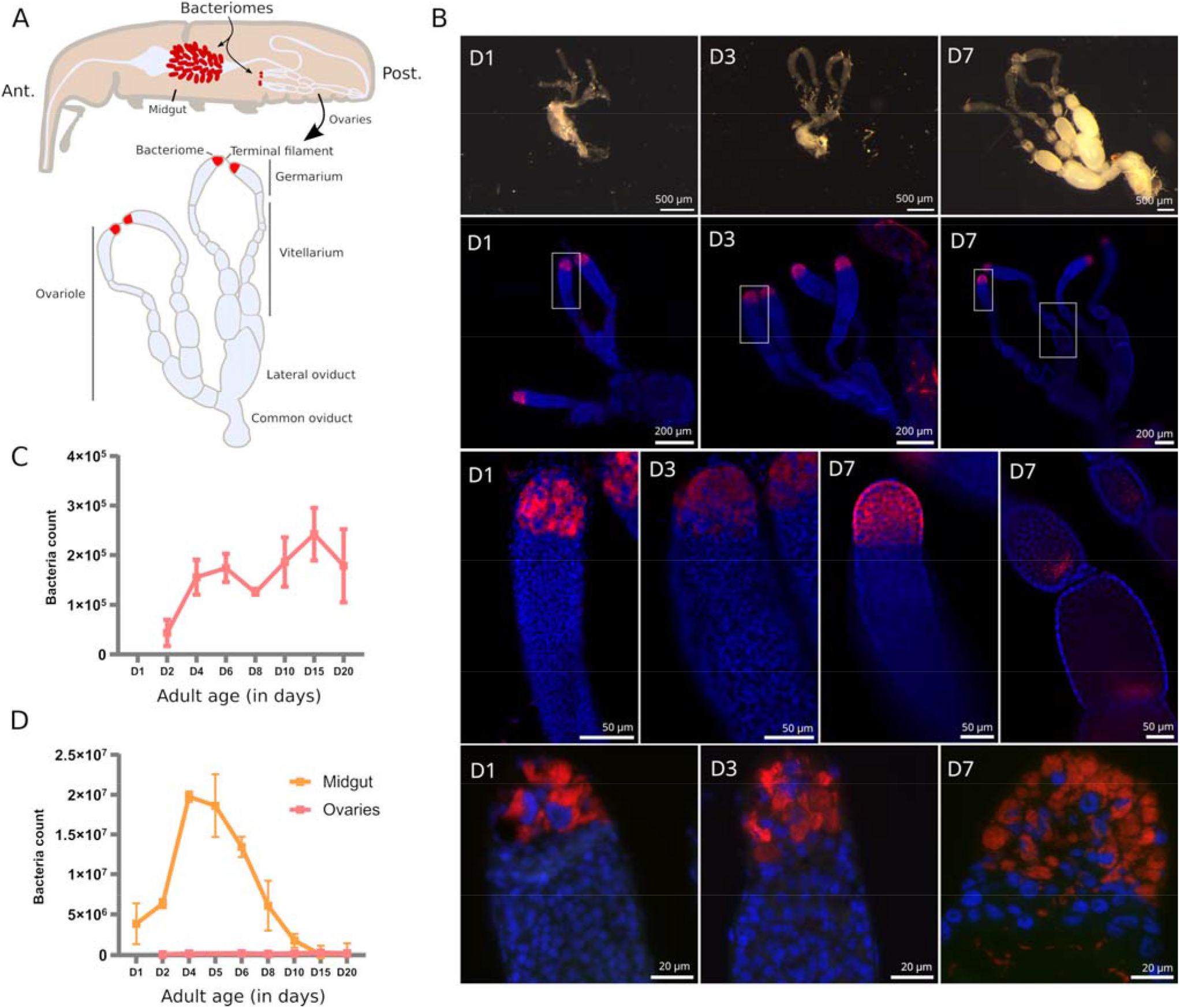
Endosymbiont localization and quantification in *S. oryzae* ovaries. A. Scheme of *S. oryzae* adult showing midgut and ovarian bacteriomes, along with a scheme of ovaries based on observation and previous reports (Perez-Mendoza et al., 2004; Nardon, 2006). *S. oryzae* harbours two ovaries with two ovarioles each, composed of a germarium, a vitellarium, lateral oviducts and common oviducts. B. Upper panel: imaging of adult female reproductive system from day-one-old females (D1) to D7. In agreement with previous observations, *S. oryzae* ovaries are fully developed in one-week-old females. Middle panel (second and third lines): whole-mount immunofluorescence against endosymbionts in D1, D3 and D7 ovaries, show the presence of bacteria in the ovarian apexes, at the bacteriomes, and also at the posterior region of oocytes (D7). Rectangles indicate the region observed in the third line. The bottom panel depicts tissue sections of germaria immunofluorescence against endosymbionts. The bottom-right image shows bacteria in the germarium, outside of the bacteriome (D7). C-D. Quantification of endosymbiont load by flow cytometry in ovaries and gut (gut from (Dell’Aglio et al., 2022)). Endosymbiont load increases in the first couple of days in both ovaries and gut bacteriomes, although there are a hundred times more in the gut. After the first week of adulthood, gut endosymbionts are lost while ovarian ones are maintained. D depicts the age in days.

Maintenance of endosymbionts across generations is often ensured by vertical transmission through the female germline (Kikuchi, 2009; Kupper et al., 2016; Maire et al., 2020a), although cases of transmission through milk-gland secretions have been described in Tsetse flies (Aksoy et al., 1997; Attardo et al., 2008). Vertical transmission of *S. pierantonius* is thought to occur during embryogenesis, with infection of primordial germ cells by bacteria (Mansour, 1930; Tiegs and Murray, 1938; Heddi et al., 1999; Heddi and Nardon, 2005). In 1930, Mansour meticulously described the female genital organs along with the ovarian bacteriomes of *S. oryzae*, and suggested that endosymbionts are able to exit the bacteriocytes and migrate toward the oocytes and nurse cells, thus infecting the developing oocytes (Mansour, 1930). However, in 2006, Nardon challenged such a hypothesis since he failed to identify bacteria migrating from the apical bacteriome to the oocytes by using autoradiography methods (Nardon, 2006). Therefore, the precise mode of transmission of *S. pierantonius* in *S. oryzae* remains unclear, along with endosymbiont regulation in ovaries.

Due to the differences in tissue localization, but also to the differences in the dynamics of bacterial populations, distinct endosymbiont regulatory mechanisms likely occur between midgut and ovaries. Here, in order to uncover potential new regulators of ovarian endosymbionts in *S. oryzae*, we provide an exploratory analysis of the first transcriptome of the bacteriome-containing germarium (Figure 1A), throughout young adult development, from one-day-old adults to sexually mature ones (day 7). Contrary to midguts, we show that the endosymbiont-specific AMP ColA is downregulated in the germarium at the onset of sexual maturity, along with several other AMPs, the transcription factor *relish* and the receptor *pgrp-lc*. Downregulation of immune effectors is concomitant with upregulation of *caudal*, a negative regulator of the IMD pathway, which is downregulated in midgut bacteriomes at the same stages. Finally, *pgrp-lb* is upregulated, independently of *caudal* repression, potentially inhibiting the re-activation of the IMD pathway. We hypothesize that downregulation of ColA and related AMPs, allows bacterial release from bacteriocytes, similar to RNA interference experiments of *colA* in the midgut (Login et al., 2011), potentially allowing oocyte infection and endosymbiont transmission.

## Methods

### Insect rearing

The *S. oryzae* population used in this study was previously sampled in the Azergues valley, France in 1984, and has been reared in a climate chamber (27.5 °C, 70% relative humidity, no light source) on wheat grains ever since. In these conditions, the timespan between egg laying and the emergence of adults from the grain is one month. Adults emerging from the grain are fully formed, although their cuticle is reinforced during the two-three days following emergence (Vigneron et al., 2014; Dell’Aglio et al., 2022), together with sexual maturation (Perez-Mendoza et al., 2004; Dell’Aglio et al., 2022). The time between the end of metamorphosis and emergence has been calculated to be around three to four days (Vigneron et al., 2014; Dell’Aglio et al., 2022). Because young adults are present within the grain, we defined Day 1 as adults inside the grain that are unable to walk and show a light cuticle, Day 3 corresponds to animals exiting the grain naturally, and Day 7 is four days after the animals exited the grain.

### Observation of ovarian development in young adults

Ovaries were dissected in buffer TA for each time point (one, three and seven-day-old females) under a stereomicroscope. Images were captured using an Olympus XC50 camera and processed with ImageJ software (v1.53t).

### Endosymbiont localization by Fluorescent *in situ* hybridization

Five ovaries per replicate were dissected in buffer TA for each time point (one, three and five-day-old females), and immediately immersed in PFA 4% in PBS. Two replicates were performed for each timepoint. Fluorescence in situ hybridization (FISH) was carried out as previously described (Maire et al., 2019).

Briefly, for whole mount observation, ovaries were rinsed, permeabilized in acetic acid 70% at 60°C for 1 minute and deproteinized in pepsine 0,1 mg/ml in hydrochloric acid 0,01N for 20 minutes at 37°C. Then prehybridization was carried out for 30 minutes at 45°C in a prehybridization buffer (79% of hybridization buffer, 20% of Denhardt (Ficoll 10%, Polyvinylpyrrolidone 10%, Bovine Serum Albumin 10%), and 1% SDS), prior to hybridization at 45°C in a hybridization buffer (NaCl 0.9 M, Tris 20 mM, EDTA 5 mM, pH 7.2) with 10μM TAMRA-labeled probe targeting *S. pierantonius* (5’-/56-TAMN/ACC-CCC-CTC-TAC-GAG-AC-3’ - Integrated DNA Technologies, Inc.). After 3 hours of incubation, samples were then washed with hybridization buffer with SDS 0,1%, rinsed in PBS and distilled water, and immersed overnight in an aqueous mounting medium with an anti-fading agent (Fluoro Gel with DABCO™) with DAPI 3μg/ml.

Samples were observed with a Leica DMi8 widefield microscope with the THUNDER imager system, and images were acquired with the K3M monochrome camera. For each sample, multistack images were acquired (two-channels with LED 405 nm and TXRed filter cube / Z stacks) and processed as described: images were first deconvolved using Thunder integrated software with the Small Volume Computational Clearing method, then, for larger views, the Z stacks were combined using maximum intensity Z projection method (Leica LasX software). For smaller views, only one stack was chosen to maximize resolution.

Concerning tissue sections, fixed samples were washed repeatedly with PBS before embedding the tissue in 1.3% agar. Next, samples were dehydrated through a graded ethanol (EtOH) series and transferred to butanol-1, at 4 °C, overnight. Samples in agar were then embedded in melted Paraplast. Tissue sections (3-μm thick) were cut using a rotary microtome (HM340E Thermo Fisher Scientific). Sections were placed on poly-lysine-coated slides, dried overnight in a 37 °C oven, and stored at 4 °C prior to further treatments. After methylcyclohexane dewaxing, sections were covered with a drop of 70% acetic acid. Deproteinization of slides was performed in hydrochloric acid 0.01 N with pepsin 0.1 mg/ml for 10 min at 37 °C. Prehybridization and hybridization were carried out as in the above whole mount experiments. Samples were observed with the Olympus IX81 epifluorescence microscope through appropriate filters for fluorescence. Images were captured with the XM10 monochrome camera and processed with ImageJ software (v1.53t).

### Flow cytometry quantification of ovarian endosymbionts

*S. oryzae* ovaries were dissected in buffer TA (35 mM Tris/HCl, 25 mM KCl, 10 mM MgCl2, 250 mM sucrose, pH 7.5) under a stereomicroscope. Three biological samples per time point were collected, each composed of 10 ovarian systems. Flow cytometry was performed exactly as described previously (Dell’Aglio et al., 2022).

### RNA extraction and sequencing

Germaria from young adults at Day 1 (D1), Day 3 (D3) and Day 7 (D7) were dissected in buffer TA (35 mM Tris/HCl, 25 mM KCl, 10 mM MgCl_2_, 250 mM sucrose, pH 7.5) in order to span the sexual maturity of adults (Maire et al., 2020b), along with the increase in endosymbiont load (Vigneron et al., 2014). Twenty to 28 ovarian apexes were dissected per triplicate. RNA extraction was performed using Ambion RNAqueous micro kit (AM1931). RNA library construction was performed at GenomEast platform, using Truseq stranded mRNA library construction from Illumina. Libraries were sequenced at the same platform, in a Hiseq 4000 sequencer with a 50 bp single-end chemistry. Around 30 to 40 M reads were obtained for each sample.

### RNA sequencing analysis

All RNAseq datasets produced in this manuscript (PRJNA918856), and previously available (PRJNA918957) have been processed with the following pipeline. Quality check was performed with FastQC v0.11.8 (Andrews, 2010), and raw reads were quality trimmed using trim_galore from Cutadapt v0.6.7 (Martin, 2011), then mapped to the *S. oryzae’s* genome (GCA_002938485.2 Soryzae_2.0) using STAR v2.7.3a (Dobin et al., 2013), yielding around 90% of reads mapped. Then, uniquely mapping reads were counted with featureCounts v2.0.1 ((Liao et al., 2014), Table S1), and mapping quality was assessed by multiqc v1.13 (Ewels et al., 2016). Differential expression analysis was performed with the AskoR package (Alves-Carvalho et al., 2021). AskoR is an R pipeline that was used to convert raw read counts to counts per million (CPM), filter lowly expressed genes (CPM <0.5) and normalize counts based on trimmed mean of M values (TMM). Lists of differentially expressed genes (DEGs) were obtained based on EdgeR (Robinson et al., 2010) likelihood ratio test (LRT). Principal component analysis (PCA) was performed based on normalized counts. Coexpression analyses were also obtained through AskoR by using the coseq package (Rau and Maugis-Rabusseau, 2018). The number of ideal clusters were chosen based on the best cluster probability. Finally, Gene Ontology (GO) (Ashburner et al., 2000; The Gene Ontology Consortium, 2021) annotations were obtained from the web version of eggNOG mapper v2 (Cantalapiedra et al., 2021) using default parameters and eggNOG 5 database (Huerta-Cepas et al., 2019). Functional enrichment was performed using clusterProfiler (Wu et al., 2021). GO terms and KEGG pathways (Kanehisa and Goto, 2000; Kanehisa, 2019; Kanehisa et al., 2023) with q-values smaller than 0.05 were defined as significantly enriched and GO terms were reduced to a set of non-redundant terms with the use of REVIGO tool (Supek et al., 2011). Specific lists of genes were used for functional enrichment analyses: genes sustaining significant expression in samples (normalized CPM > 100 in all three D1, D3, and D7 RNAseq from ovarian germarium yielding 1568 genes) were chosen based on the distribution of CPM counts in all three biological replicates (Figure S1). Annotations for immune-related genes were taken from Additional File 1 from (Parisot et al., 2021). Most graphic outputs are either performed by AskoR or in R, using ggplot2 (Wickham, 2009).

## Results and Discussion

### Endosymbiont localization and load in ovaries of young female adults

*S. oryzae* female reproductive system is composed of two telotrophic meroistic ovaries, as nurse cells are present within the germarium and connected to oocytes by nutritive cords (described in (Tiegs and Murray, 1938; Perez-Mendoza et al., 2004; Nardon, 2006) and depicted in Figure 1A). Each ovary is composed of two ovarioles connected by a lateral oviduct merging into a common oviduct. Oogenesis is progressive from the germarium, through the vitellarium, and finishes at the lateral oviduct (Tiegs and Murray, 1938; Nardon, 2006). Fertilization happens thanks to the spermatheca glands and the bursa copulatrix (Tiegs and Murray, 1938; Perez-Mendoza et al., 2004).

In young pre-emerged females (D1), and newly emerged ones (D3), no oocytes can be distinguished, and the germarium and vitellarium are confounded (Figure 1B, upper panel). Nevertheless, the bacteriome is clearly visible from D1 onwards, in agreement with their observation as early as in larvae sections (Maire et al., 2020a). From one-week-old females (D7) the ovaries harbour mature oocytes, and the germarium is clearly visible along with the apexes containing the bacteriomes (Figure 1B, upper panel), in agreement with previous observations (Perez-Mendoza et al., 2004). Sexual maturation of young females is therefore achieved around one-week-old females (Perez-Mendoza et al., 2004; Campbell, 2005).

In adult ovaries, *S. pierantonius* was described within the bacteriome and germ cells (oocytes and nurse cells) (Tiegs and Murray, 1938; Nardon et al., 1998; Heddi et al., 1999). At early stages (D1), individual bacteriocytes can be clearly distinguished in the ovarian apexes, while at later stages, upon sexual maturation (D7), an important mass of bacteriocytes can be observed, suggesting an increase in endosymbiont load (Figure 1B, middle panels). Endosymbionts are present in ovarian bacteriomes but at later stages (D7) can also be frequently observed in the germarium (Figure 1B, bottom panel). This observation reinforces the hypothesis that endosymbionts might be able to infect oocytes *via* the ovarian bacteriomes. Whether endosymbionts are able to migrate as previously observed in the gut during metamorphosis (Maire et al., 2020b), or if infected nurse cells are able to carry endosymbionts that infect the oocytes, remains to be defined. As described by Tiegs and Murray in 1938, (Tiegs and Murray, 1938), and observed here, in oocytes, endosymbionts are present in the posterior pole, where primordial germ cells will be present, and the formation of “germline” bacteriomes could occur (Figure 1B, middle panel).

Quantification of bacteria in whole ovaries shows a five-fold increase in load from young adults to sexually mature ones (Figure 1C). The load of endosymbionts in mature ovaries remains significantly smaller than in adult midguts, even at early stages (Figure 1D), demonstrating the extreme amplification of midgut endosymbionts during the first week of adulthood. Given the different dynamics and density of endosymbionts between the midgut and ovarian bacteriomes, we investigated if the immune pathways associated with endosymbiont control in *S. oryzae* are accordingly modulated.

### Transcriptional landscape of *S. oryzae* germarium

The germarium, containing the bacteriomes, was carefully dissected from one (D1), three (D3), and seven-day-old adults (D7), the latter which corresponds to *S. oryzae* sexual maturity and mild increase in endosymbiont load, and submitted to RNA extraction and sequencing (Table S1). Principal component analysis of mapped reads shows a clear difference in transcriptome landscapes between D1, D3 and D7 (Figure 2A) suggesting time point-specific transcriptome signatures. In order to understand the overall common landscape of ovarian bacteriomes, genes commonly expressed in the three-time points were subjected to gene ontology (GO) enrichment (Figure 2B, genes are considered expressed when normalized CPM > 100 in all time points as explained in the Material and Methods and Figure S1). Among the most enriched GO terms, we detected typical ovary-developmental and cell division terms such as “germarium-derived egg chamber formation”, “morphogenesis of follicular epithelium”, and “meiotic cell cycle”, which validate the tissuespecificity of our RNAseq approach. Furthermore, several other terms related to developmental functions and regulation of gene expression were also commonly enriched (Table S1). KEGG pathway analysis showed significant enrichment in translational activity “Ribosome”, “Protein processing in endoplasmic reticulum”, and “Proteasome” (Figure 2C), as previously observed through histochemical analysis of nurse cell cytoplasms (Nardon, 2006). Finally, terms related to signaling and response to stimuli, including “regulation of symbiosis, encompassing mutualism through parasitism”, “interspecies interaction between organisms”, and finally, “immune effector process” were also enriched in the shared germarium transcriptomes.

**Figure 2.**
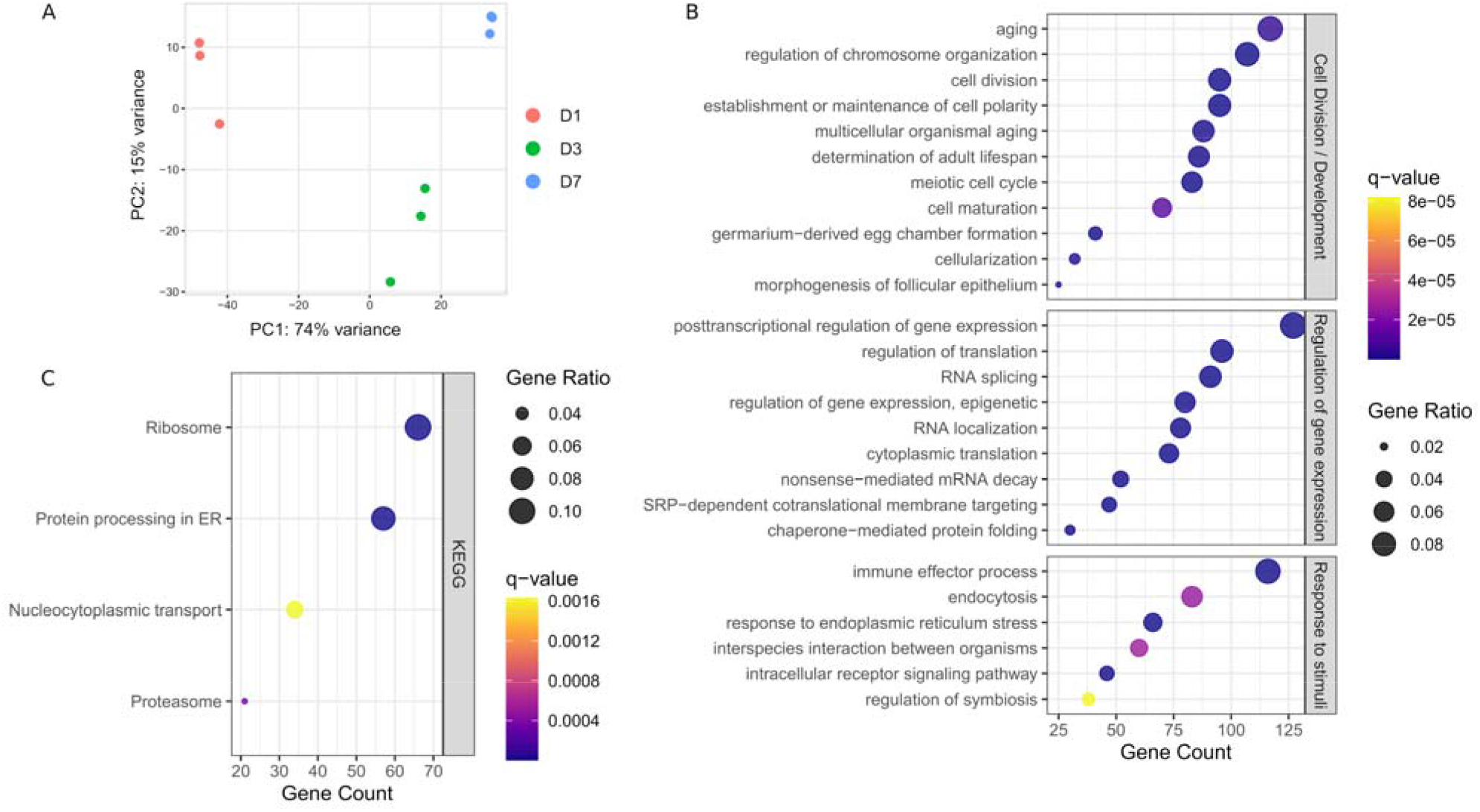
Global transcriptome analysis of *S. oryzae* germarium. A) Principal component analysis (PCA) of normalized gene counts at day-one, -three and -seven (D1, D3, and D7) show distinct transcriptome signatures. B) Biological process terms from gene ontology (GO) enrichment analysis on genes with normalized counts per million (CPM) > 100 in all three developmental times. C. KEGG pathway enrichment analysis on genes with normalized counts per million (CPM) > 100 in all three developmental times. See Material and Methods and Figure S1 for more information on CPM filtering.

### High transcriptional dynamics across young adult germarium

There were 3 829 genes differentially expressed between D1 and D3, and 2 350 between D3 and D7 (Figure 3A, adjusted p-value < 0.05 & |log2 fold change| > 1), confirming the specific transcriptomic signature of each time-point observed in Figure 2A. While between D7 and D3 most genes are upregulated, in the early stages of adulthood (D1 and D3), differentially expressed genes are more contrasted, albeit a majority are upregulated (Figure 3B). In order to assess the overall transcriptional dynamics we performed a co-expression analysis using AskoR (Alves-Carvalho et al., 2021) based on the coseq package (Rau et al., 2023). The most well-suited number of clusters, based on probability analysis and redundancy of overall cluster expression profile was two, with one cluster grouping 1 721 genes progressively upregulated from D1 to D7, and the mirror cluster, harbouring 3 240 genes with a decrease in expression from D1 to D7 (Figure 3C and Figure S2-3).

**Figure 3.**
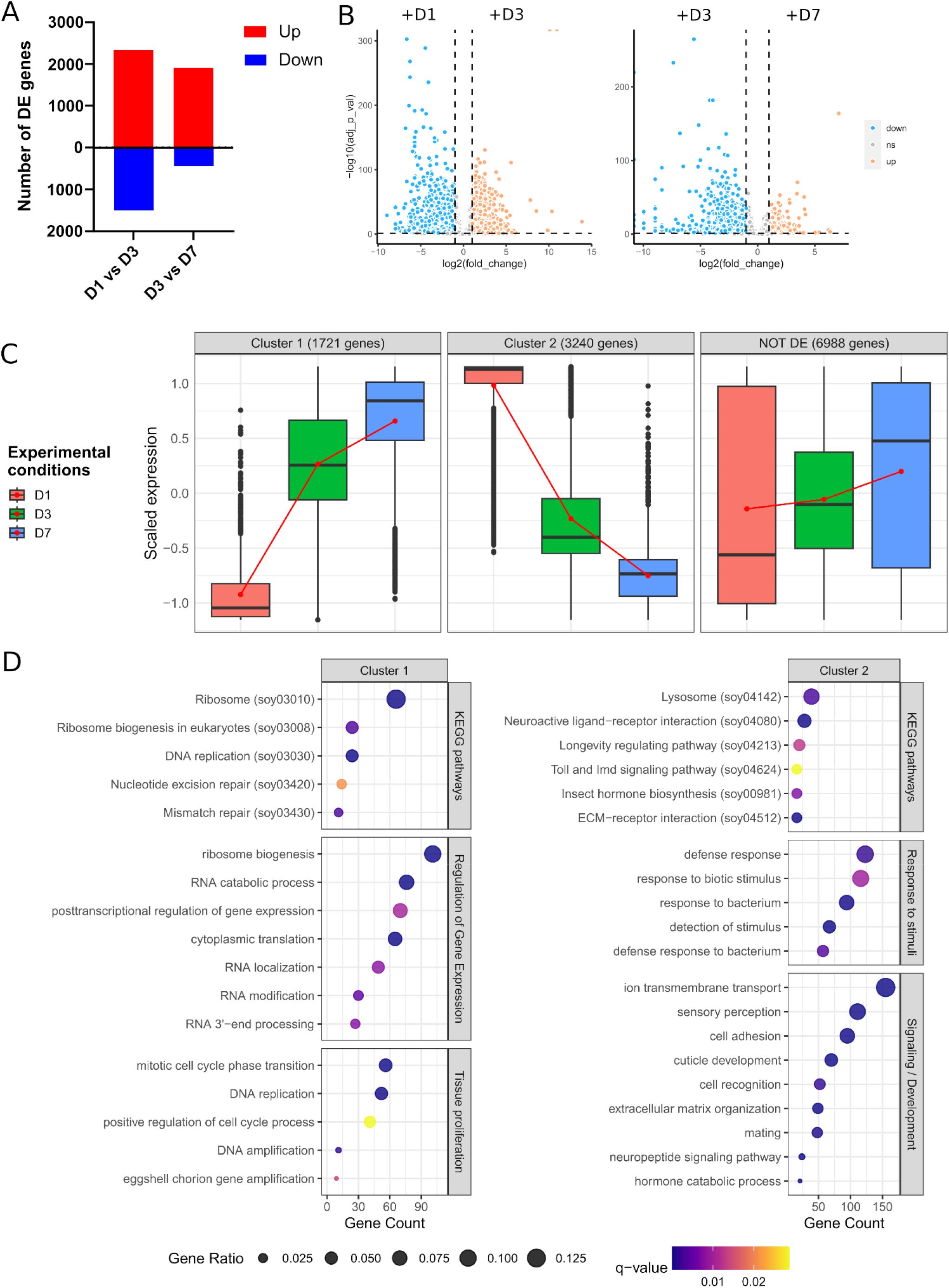
Differential expression analysis across germarium development. A. Number of genes up- or down-regulated between D1 and D3, and D3 and D7. B. Volcano plots depicting the overall differences in gene expression between the conditions tested. C. Clustering of gene expression throughout ovarian development. Two clusters are depicted, with gene expression progressively increasing between early and late females (cluster 1), or genes down-regulated from D1 to D7. D. GO and KEGG pathway enrichment analysis of genes belonging to clusters 1 and 2. Genes non-differentially expressed can be found in Figure S4.

Functional analysis of genes increasingly upregulated from D1 to D7 (cluster 1) depicted an enrichment in development, cell proliferation and regulation of gene expression, with the most significant terms being related to translation and ribosome biogenesis (Figure 3D), attesting to the high translational activity during germarium development as described previously (Nardon, 2006), and observed in the overall germarium transcriptome (Figure 2C). A similar analysis performed on cluster 2, i.e., genes progressively downregulated from D1 to D7, showed different enrichment terms, the most significant ones related to signaling, sensory perception, extracellular matrix (ECM) and neuroactive ligand-receptor interaction (Figure 3D). Indeed, neuropeptides are known in insects to regulate most physiological and developmental processes, including the maturation and functioning of female and male reproductive organs (Gäde and Hoffmann, 2005). Similarly, the roles of ECM in the ovary have been well documented in other species, including *Drosophila* (Rodgers et al., 2003; Woodruff and Shea, 2007; Luo et al., 2021). Other detected enriched terms were related to response to bacteria, “Lysosome” and “Toll and IMD signaling pathway”. The IMD pathway is indeed involved in *S. pierantonius* regulation (Maire et al., 2018), suggesting a decrease in immune activity from D1 to D7 bacteriomes. Lysosomes, on the other hand, are associated with autophagy, which has been linked to germline maintenance in *Drosophila melanogaster* (Nezis et al., 2009). In addition, midgut endosymbiont recycling in *S. oryzae* young adults is also a process governed by apoptosis and autophagy (Vigneron et al., 2014). In ovarian bacteriomes, bacterial cell lysis has been observed through autoradiography (Nardon, 2006), suggesting ovarian endosymbionts might suffer a tight control of population size. It is important to note that endosymbiont population size in ovarian bacteriomes is genetically controlled as elegantly shown by two different approaches: in females where ovaries contained a single ovariole instead of two, bacteriomes could be present twice, harbouring twice as much bacteria as an ovariole containing just one bacteriome (Grenier and Nardon, 1994); in addition, when crossing animals containing different loads of ovarian endosymbionts, Nardon and colleagues hypothesized that a genetic factor governs endosymbiont population numbers (Nardon et al., 1998). Therefore, it is tempting to suggest that autophagy mechanisms could be implicated in the regulation of the endosymbiont population in the ovaries of *S. oryzae*. Curiously, while autophagy-related genes are evenly expressed throughout the germarium development studied here (Figure S4), lysosome enrichment is only observed in cluster 2 genes, i.e., genes expressed at D1 and progressively downregulated up to D7 (Figure 3D).

Finally, as a comparison, non-differentially expressed genes (or genes commonly expressed in all stages) recapitulate some of the GO terms obtained when looking at the global transcriptome of the germarium, for instance, “multi-organism reproductive process” (Figure S4). Collectively, the functional enrichment analysis suggests that endosymbiont control might differ in immature ovaries compared to fully developed ones.

### Immune responses against *S. pierantonius*

The clustering coupled with GO terms and KEGG pathway enrichment analysis suggested a potential relaxation of immune pathways from D1 to D7 in ovarian apexes, despite an increasing endosymbiont load in this tissue over these time points (Figure 1C). To better understand the immune regulation in germaria, we used a gene-list-centered analysis, based on previous manual annotation of immune-associated genes in the *S. oryzae* assembly (Parisot et al., 2021). In order to uncover the specificity of ovarian bacteriome regulation, the expression of these immunity-related genes has also been verified in previously available RNAseq data from midguts of D1, D3 and D7 adults (PRJNA918957). This comparison not only allowed us to detect tissue-specific immune-related genes but also to uncover novel endosymbiont regulators.

Out of the 20 manually annotated immune effectors in the *S. oryzae’*s genome (Parisot et al., 2021), we detected 13 expressed in at least one germarium stage, including 11 AMPs (Figure 4A and Figure S5A). Among those, six have already been described as up-regulated in the gut-associated bacteriome at larval stages upon bacterial or TCT-challenge: *luxuorisin (lux), glycine-rich AMP like (gly-rich-AMP), sarcotoxin (srx), diptericin-2 (dpt-2), defensin, colA* (Masson et al., 2015b; Maire et al., 2018; Ferrarini et al., 2022). Furthermore, *colA* is expressed under non-challenged, standard conditions of endosymbiosis in the gut-associated bacteriome, where it was shown to act as an important endosymbiont regulator (Login et al., 2011). The germarium transcriptomic data are in line with a previous description of ColA expression in ovaries, using immunostaining (Login et al., 2011), but further reveal that *colA* is not the most expressed AMP gene in the ovarian-associated bacteriome, as *dpt-2*, *dpt-like partial* and *gly-rich-AMP* are up to ~500 times more expressed than *colA* in a given time-point (Figure 4 and Figure S5). The expression of 13 immune effectors in the germarium indicates an immune protection of this organ. These immune effectors could be involved in endosymbiont regulation and/or immune protection of the ovaries from non-symbiotic bacteria. In contrast, in the available transcriptome data of midguts, containing the bacteriomes present at the midgut caeca, only nine genes annotated as immune effectors are expressed from D1 to D7, including all those expressed in the germarium (Figure 4B, Figure S5B). This could suggest a stronger control of endosymbiont proliferation in the germarium in the early beginning of adult life.

**Figure 4.**
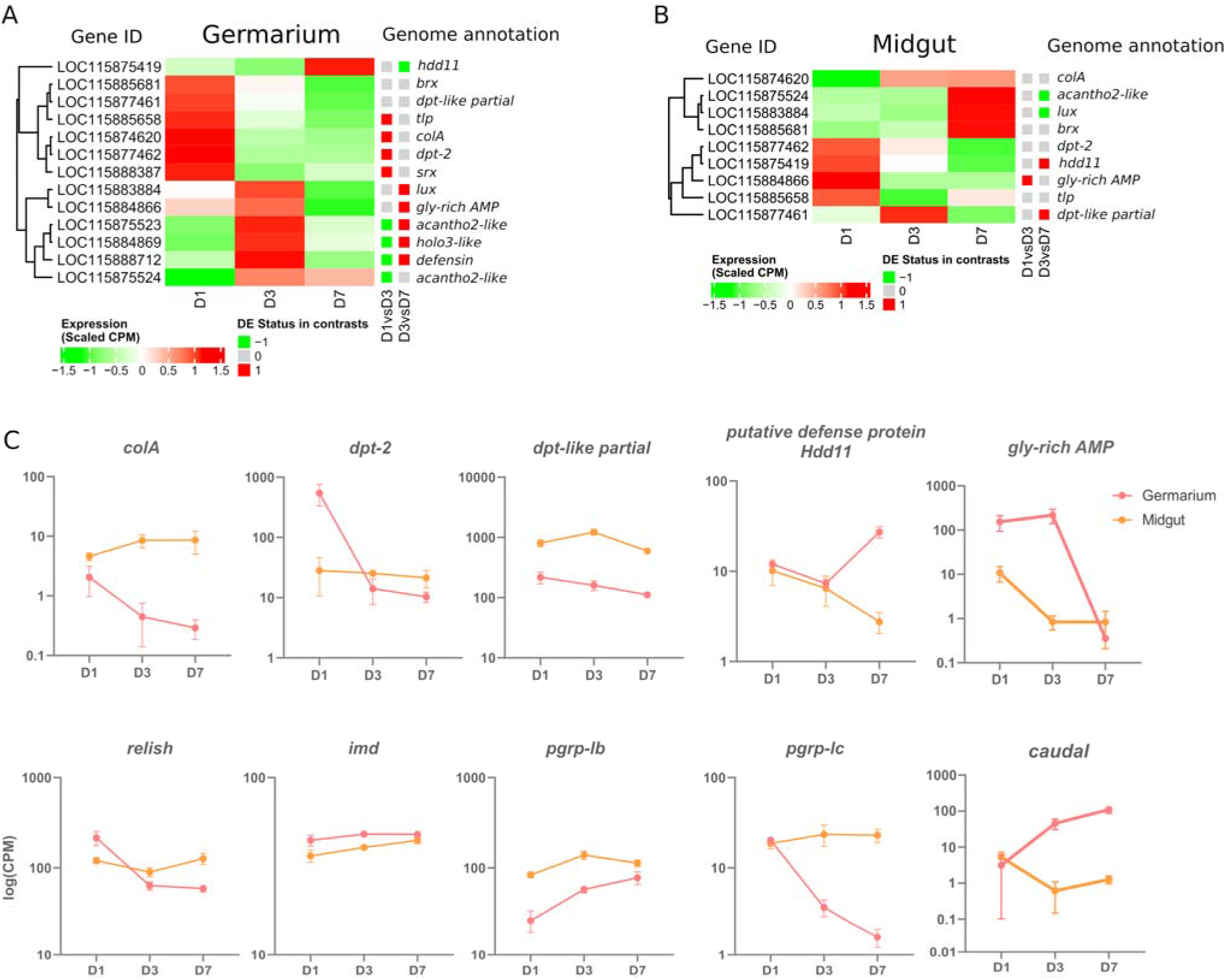
Immune-gene-centered analysis in the germarium and midguts of D1, D3 and D7 adults. A. Heatmap of scaled CPM expression of AMPs in germarium (A) and midgut (B). Differentially expressed status between D1 and D3, and D3 and D7 is depicted with a coloured squared (red upregulation, green downregulation). C. Examples of genes expressed at specific time points in midgut and germarium. Log of Normalized CPM are depicted.

Curiously, immune effector expression is time-dependent with most genes significantly upregulated at D1 with a progressive downregulation up to D7 (*colA, dpt-2, thaumatin-like protein* (*tlp*) and *sarcotoxin* (*srx*), Figure 4A and S5A). Other AMPs are upregulated at D1-D3 or only at D3, such as *acanthoscurrin-2-like (acantho2-like), holotricin-3-like (holo3-like), gly-rich-AMP, defensin* and *lux*. The AMP *acantho2-like* is the only one to follow endosymbiont load dynamics, increasing at D3 and remaining stable at D7 (Figure S5A). Finally, only one AMP is upregulated at D7, a *putative defense protein Hdd11*. Intriguingly, while *colA* follows the bacterial dynamics in midgut caeca (Figure 1D and 4C), we observe, in the germarium, a decrease in the expression of this endosymbiont-specific regulator, anti-paralleling the bacterial load that slowly increases in the ovaries. Hence the germarium transcriptome indicates a relatively strong expression of effectors at D1, and a controlled, down-regulated immune response at later stages. Finally, the data highlights that co-expression of AMPs are different in the ovarian and gut tissues (Figure 4A-B and Figure S5). The absence of common coexpression patterns suggests that distinct, tissue-dependent mechanisms are involved in the regulation of these immune effectors.

Among the 13 immune effectors expressed in the germarium, six have been demonstrated as regulated by the IMD pathway (*colA*, *srx*; (Maire et al., 2018, 2019), or are expected to be as they are significantly induced upon immune challenge with Gram-negative bacteria or their TCT fragments (*lux, gly-rich-AMP, dpt-2, defensin;* (Masson et al., 2015b; Ferrarini et al., 2022)). The IMD pathway (Figure 5) is known to be conserved, complete (26 annotated genes in the *S. oryzae* genome (Parisot et al., 2021)), and functional (Maire et al., 2018) in *S. oryzae*. We observe the expression of all the pathway’s genes in the germarium (Figure S6A) and midguts (Figure S6B). The immune effectors’ expression pattern in the germarium is relatively high at D1 and largely down-regulated by D7, hence strongly suggesting that the IMD pathway is activated at D1, before being tightly down-regulated, especially by D7, as observed in the functional analysis of cluster 2 genes (Figure 3D). The expression pattern of *relish*, the transcription factor of this immune pathway, also follows a decrease from D1 to D3-D7 (Figure 4C), which is in line with the fact that *relish* has been described to be up-regulated upon IMD pathway activation, contrary to *imd*, which presents a steady expression profile after immune challenge (Maire et al., 2018), and through germarium developmental stages (Figure 4C). In contrast, in midgut samples, *relish* expression follows the adult bacterial load and increases between D1 and D7 (Figure 4C). Hence, the data suggest that a tissue-specific regulation of the IMD pathway occurs in ovarian bacteriomes.

**Figure 5.**
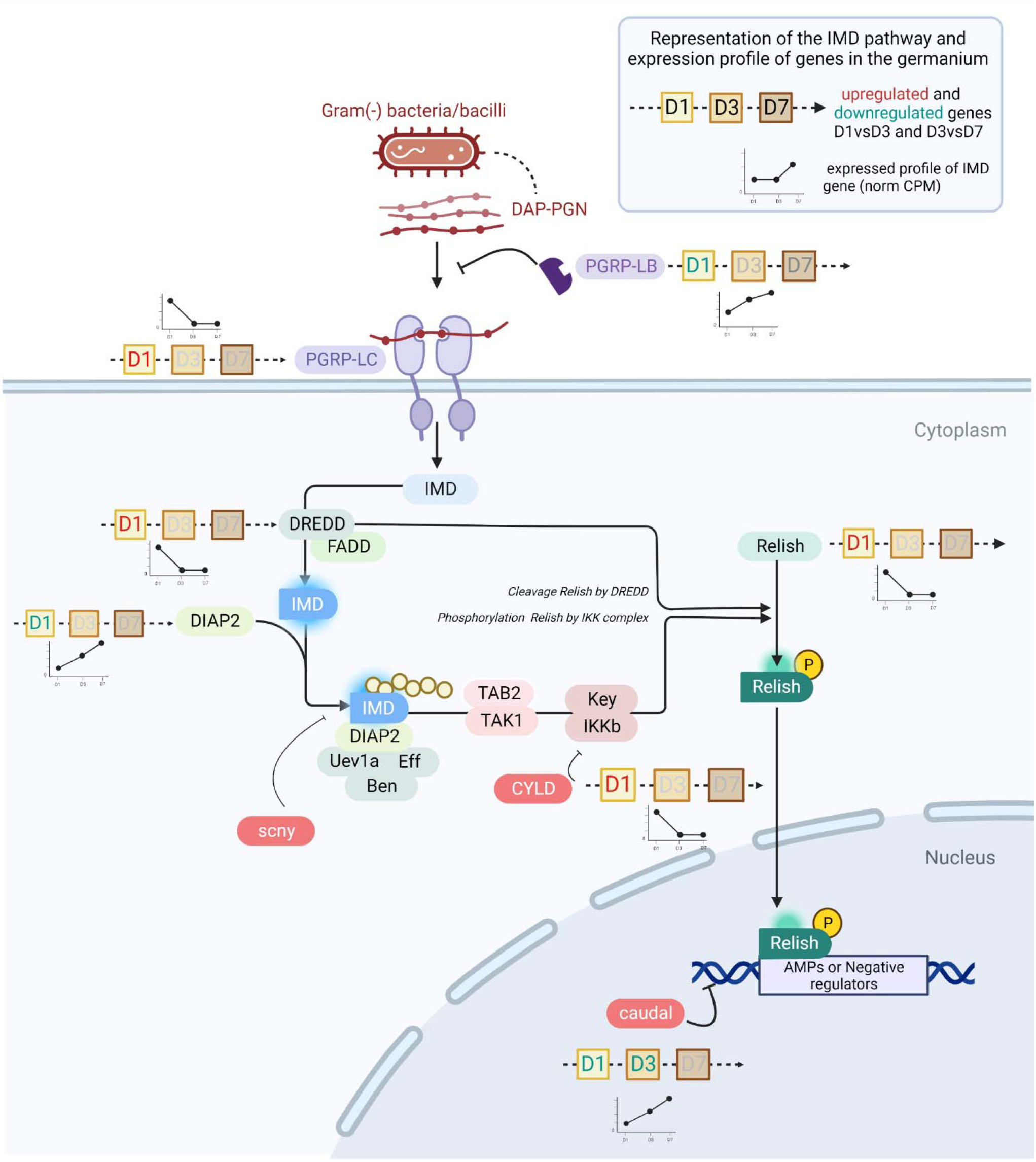
The IMD pathway in *S. oryzae* during germarium development. The expression profile of genes is depicted by line histograms and based on Figure 4. The time points studied are depicted as squares, with D1, D3 or D7 written inside. Red letters correspond to the upregulation of the gene at that stage, and greenblue letters depict downregulation. In pre-emerged females, the IMD pathway is active and AMPs are abundantly produced. In one-week-old females, *caudal* is upregulated and consequently represses *relish* expression, potentially causing downregulation of AMPs. Finally, *pgrp-lb* is constantly expressed across time points, insuring cleavage of monomeric TCT and avoiding reactivation of the IMD pathway by PGRP-LC.

Many negative regulators of the IMD pathway have been characterized in other insects, especially the genetic model *Drosophila melanogaster* (Zhang et al., 2021; Cammarata-Mouchtouris et al., 2022). Analysis of their expression patterns between D1 and D7 in the germarium suggests the potential implication of three main regulatory mechanisms in this tissue. First, we observe that the expression of *caudal* (*cad*) is up-regulated from D1 to D3 and from D3 to D7 in the ovaries but not in the gut (Figure 4C and Figure 5). *cad* is an important downregulator of the IMD pathway in presence of gut commensal bacteria in *D. melanogaster*, and acts at the most basal level of the pathway, *i.e*. by counteracting Relish effects on its transcriptional targets (Ryu et al., 2008). Interestingly, while *cad* has been shown to be involved in AMP downregulation in *D. melanogaster*, *pgrp-lb* expression was not affected (Ryu et al., 2008), a gene known to be under the control of the IMD pathway and encoding another down-regulator of the IMD pathway (Zaidman-Rémy et al., 2006). This is in line with the expression patterns observed here: while most immune effectors and *relish* expression are downregulated at D7, *pgrp-lb* expression increases from D1 to D7 (Figure 4C). Hence, by D3 and D7, PGRP-LB is likely contributing to the control of the IMD activation, by cleaving the immune elicitors of the IMD pathway in small fragments that are no longer recognized by PGRP-LC, the receptor upstream of the adaptor protein Imd (Zaidman-Rémy et al., 2006; Maire et al., 2019) (Figure 5). In *S. oryzae*, we have shown that this regulation is important for the symbiosis homeostasis by avoiding a systemic activation of the larval immune system in response to the endosymbiotic presence in the bacteriome (Maire et al., 2019). The expression of *pgrp-lb* in the germarium indicates an active regulation of the IMD pathway in this tissue, through the degradation of the TCT produced by the endosymbionts. Finally, a third regulatory mechanism is likely involved in this tissue, which relies on the downregulation of the expression of the *pgrp-lc* receptor itself (Figure 4C). *pgrp-lc* expression decreases progressively and significantly between D1 and D7 in germarium samples (Figure 4C and Figure S7). While *pgrp-lb* increases in expression with the increase in bacterial load both in midgut and in ovaries, the expression of the two other regulators is contrasted between the midgut and the ovaries: *pgrp-lc* maintains a constant expression level in midguts, and *cad* expression significantly decreases (Figure 4C).

## Conclusion

In this work, we reveal the presence of endosymbionts within the germarium, outside of bacteriomes, suggesting that bacteria might be able to actively migrate from the bacteriome apexes to the nearby vitellarium. Nevertheless, we lack to demonstrate the mechanisms associated with endosymbiont infection of primordial germ cells, and a complete analysis of embryogenesis along with the early formation of ovaries should elucidate such a question.

Regulation of the immune effectors’ expression in the germarium shows a consequent immunocompetency at D1, and a timely downregulation of the IMD pathway by D7, a stage which corresponds to the physiological (*e.g*. with cuticle completion) and sexual maturation of the cereal weevil. The immune activity at D1 could be required to either protect the ovaries from infection with opportunistic non-symbiotic bacteria, and/or to control the endosymbiont load, which remains very small compared to the population size observed in the midgut concomitantly. Subsequent downregulation of most of the immune effectors could be essential to prevent endosymbionts from being eliminated from the tissue, hence allowing their maintenance and transmission to the next generation. Not only this downregulation could be important to preserve the endosymbionts in the germarium, but to allow them to infect the growing oocytes. Artificial downregulation of *colA* expression by RNAi led to an escape of endosymbionts from the larval midgut-associated bacteriome (Login et al., 2011). Therefore, the downregulation of *colA* and other AMPs in the germarium could allow the transmission of endosymbionts to the next generation. Last, the adjustment of the immune activation along ovary maturation is likely a trade-off between the advantages of protection against pathogens and/or endosymbiont control, and the cost of the immune response itself. It is intriguing that our previous analysis of the function of *pgrp-lb* in *S. oryzae* using systemic RNAi has revealed both a systemic activation of the immune system and a delay in egg production (Maire et al., 2019). In light of these last data, we cannot exclude that the effects observed could be partially due to the local downregulation of *pgrp-lb* in the ovaries.

Finally, while this study remains descriptive, the numerous candidate genes of endosymbiont regulation attest to the importance of genome-wide transcriptomic studies. In addition to *colA*, we pinpointed *dpt-2* and *dpt-like partial* as potential immune effectors against endosymbionts, both in ovaries and midguts. The only AMP following the bacterial dynamics in ovaries, *acantho2-like*, is certainly a potential target for future functional analysis.

## Author Contributions

RR conceived the study with the help from AH and NP; AV produced the next-generation sequencing data; EDA performed the flow cytometry analysis, MGF analyzed the sequencing data; MD and SB performed the immunofluorescence and imaging analysis; AZR, JM and CVM analyzed and wrote the immune-related data analysis, RR wrote the article and all authors read, substantially edited, and approved the final draft.

## Acknowledgments

We would also like to warmly thank Sandrine Hugues and Benjamin Gillet for their efforts in producing the preliminary data that inspired this work. We would like to thank Stéphanie Robin and Fabrice Legeai for the prompt help and for teaching RR to use AskoR. RNA library construction and sequencing were performed at the GenomEast platform.

## Data Availability Statement

The datasets generated and analyzed for this study can be found at the following BioProjects PRJNA918856 (http://www.ncbi.nlm.nih.gov/bioproject/918856) for germarium RNAseq data, and PRJNA918957 (http://www.ncbi.nlm.nih.gov/bioproject/918957) for midgut RNAseq.

