## Supplementary Figures for "Transcriptional survey of ovarian bacteriomes in the cereal weevil, *Sitophilus oryzae*, shows down-regulation of immune effectors at the onset of sexual maturity"

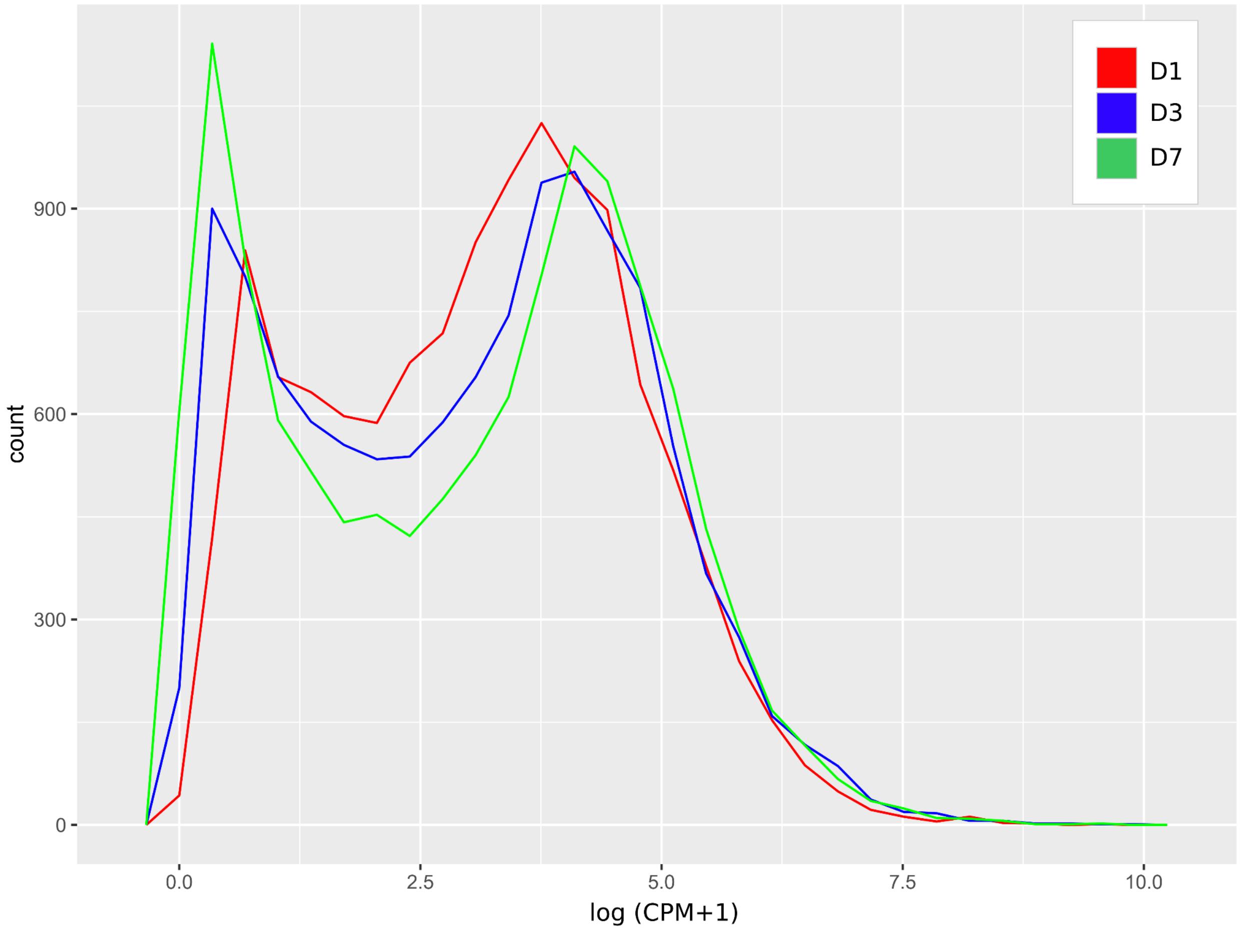


Figure S1. Distribution of log (normalized meanCPM+1) counts in the three biological replicates D1, D3 and D7. We can see that a threshold of 100 normalized CPM, corresponding to ~2, separates the large peak of lowly expressed genes from the peak of expressed genes.

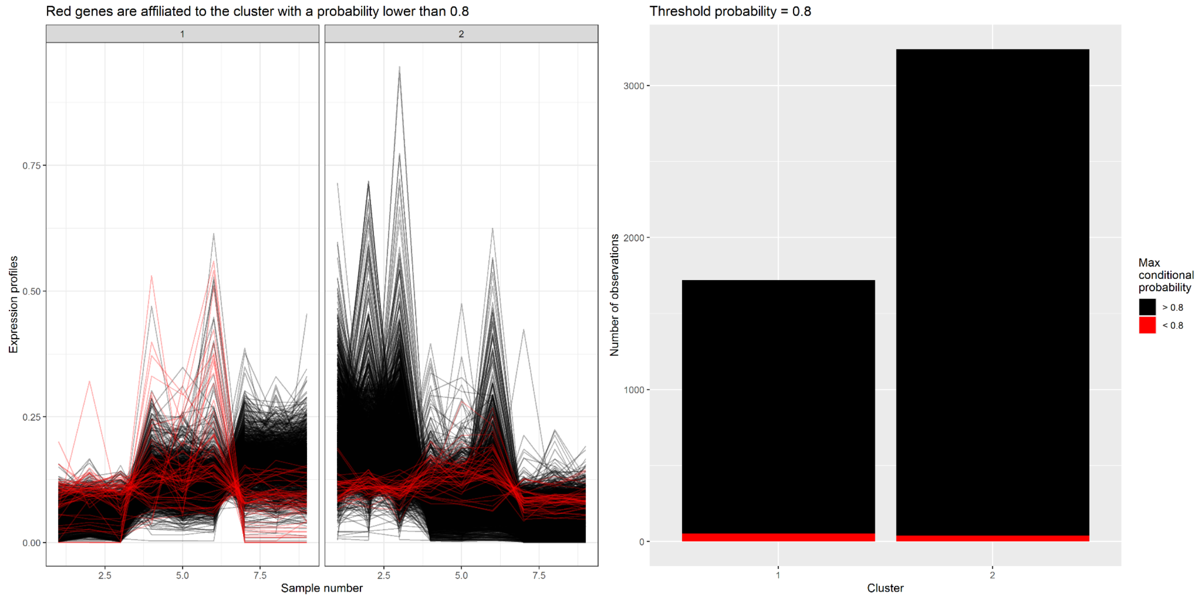


Figure S2. Probability of genes belonging to the two clusters depicted in Figure 3C

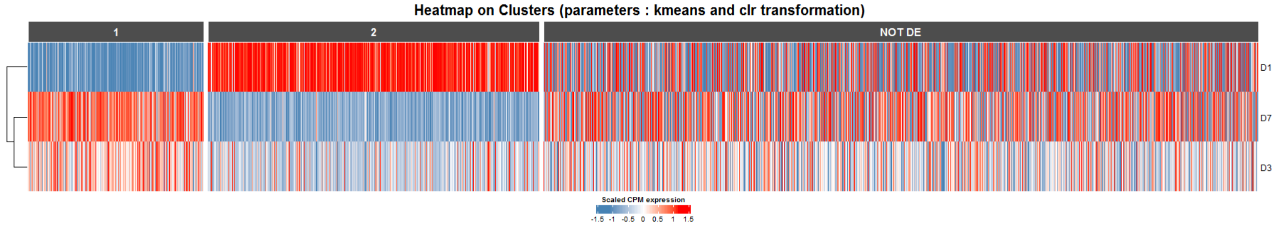


Figure S3. Heatmap of scaled normalized counts per million (CPM) expression of genes belonging to cluster 1, cluster 2 and the non-differentially expressed genes.

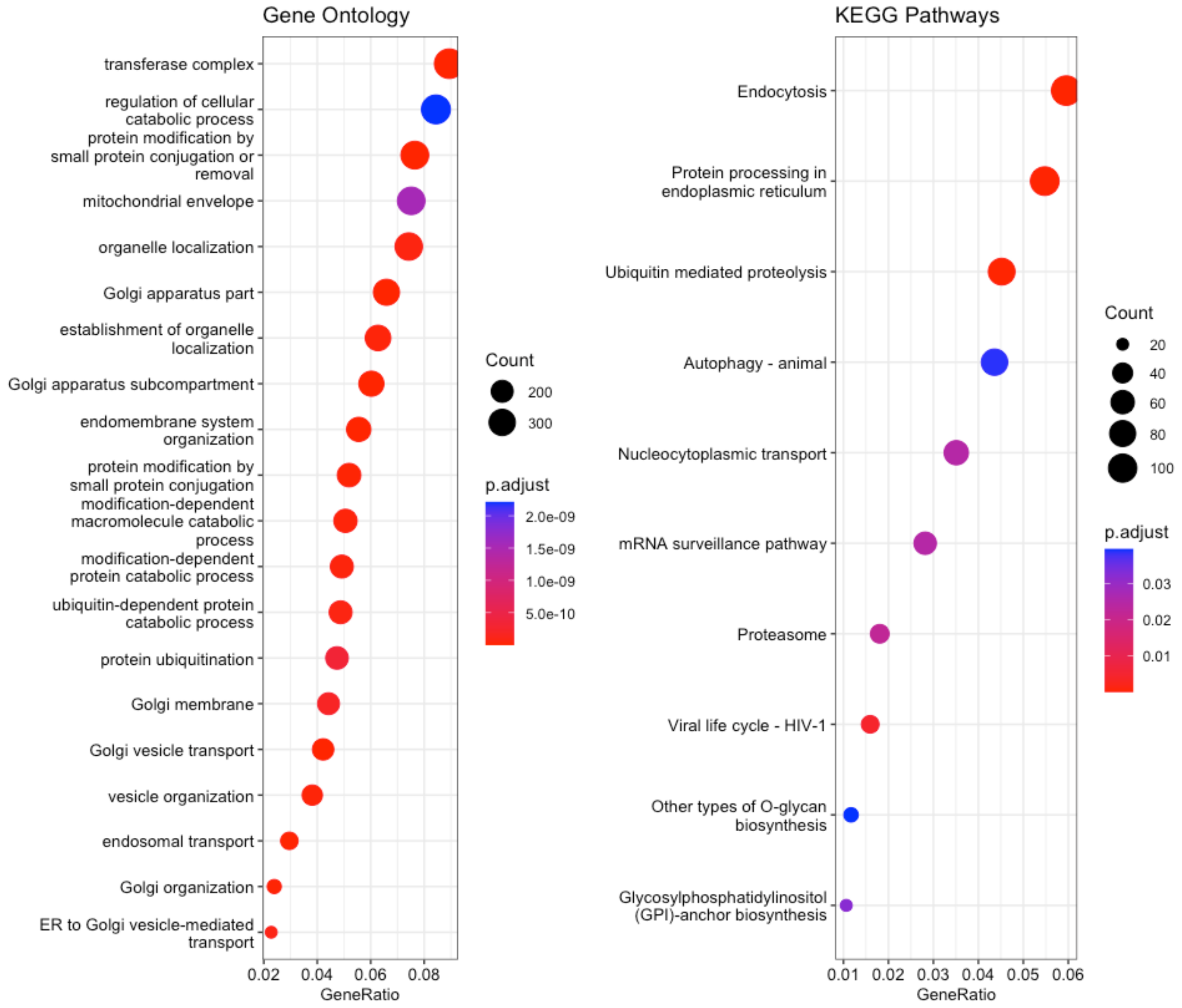


Figure S4. GO enrichment analysis of genes non-differentially expressed between conditions.

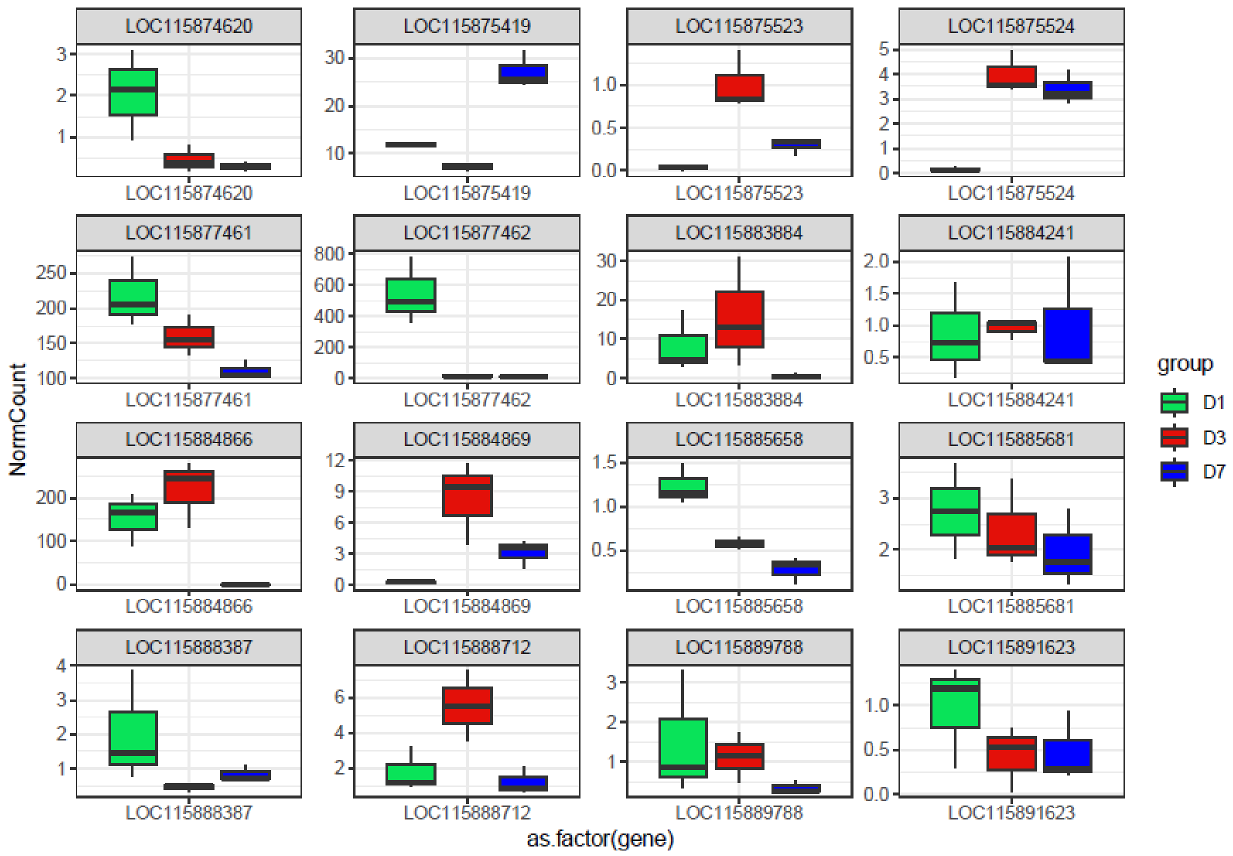


Figure S5A. Normalized counts per million (CPM) of expressed AMPs in D1, D3 and D7 germarium.

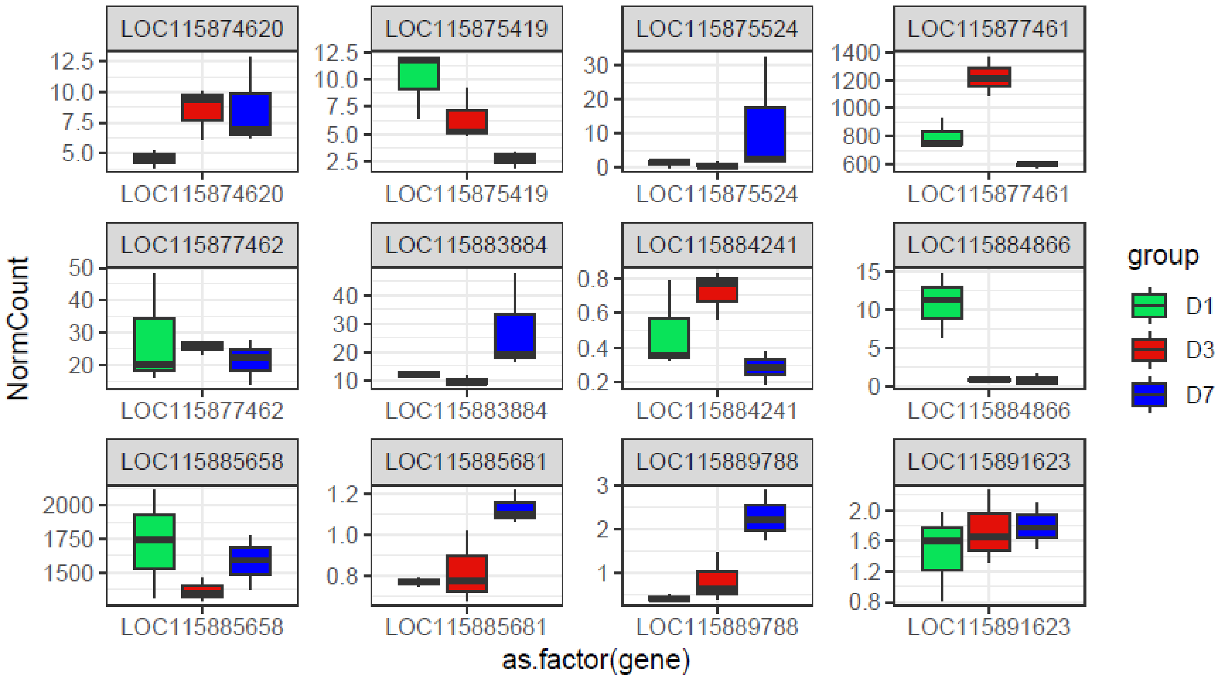


Figure S5B. Normalized counts per million (CPM) of expressed AMPs in D1, D3 and D7 midguts.

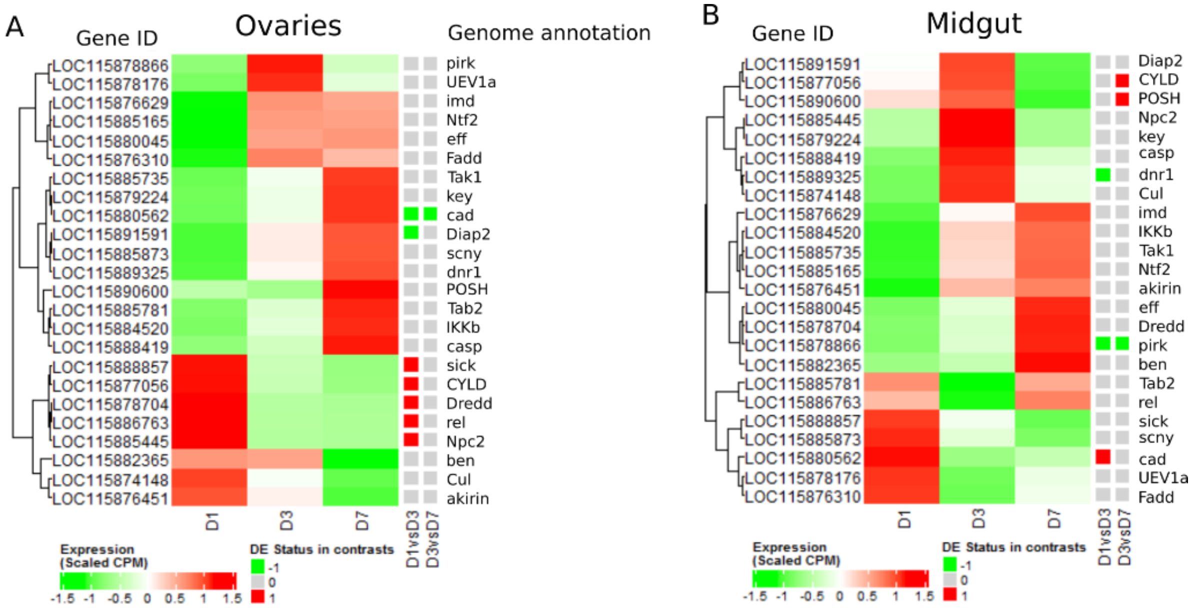


Figure S6. IMD-gene-centered analysis in the germarium and midguts of D1, D3 and D7 adults. A-B. Heatmap of scaled CPM expression of genes in germarium (A) and midgut (B).

Differentially expressed status between D1 and D3, and D3 and D7 is depicted with a coloured squared (red upregulation, green downregulation).

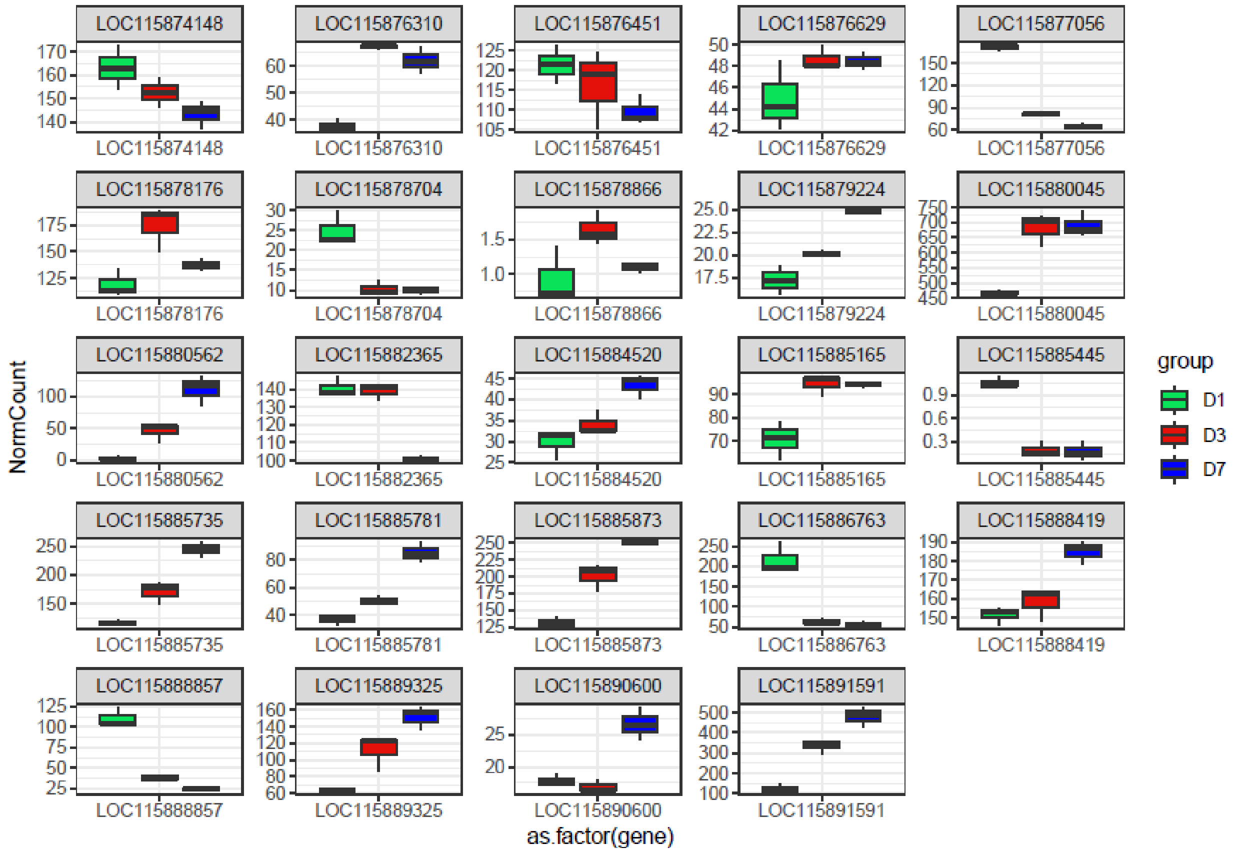


Figure S6C Normalized counts per million (CPM) of expressed IMD related genes in D1, D3 and D7 germarium.

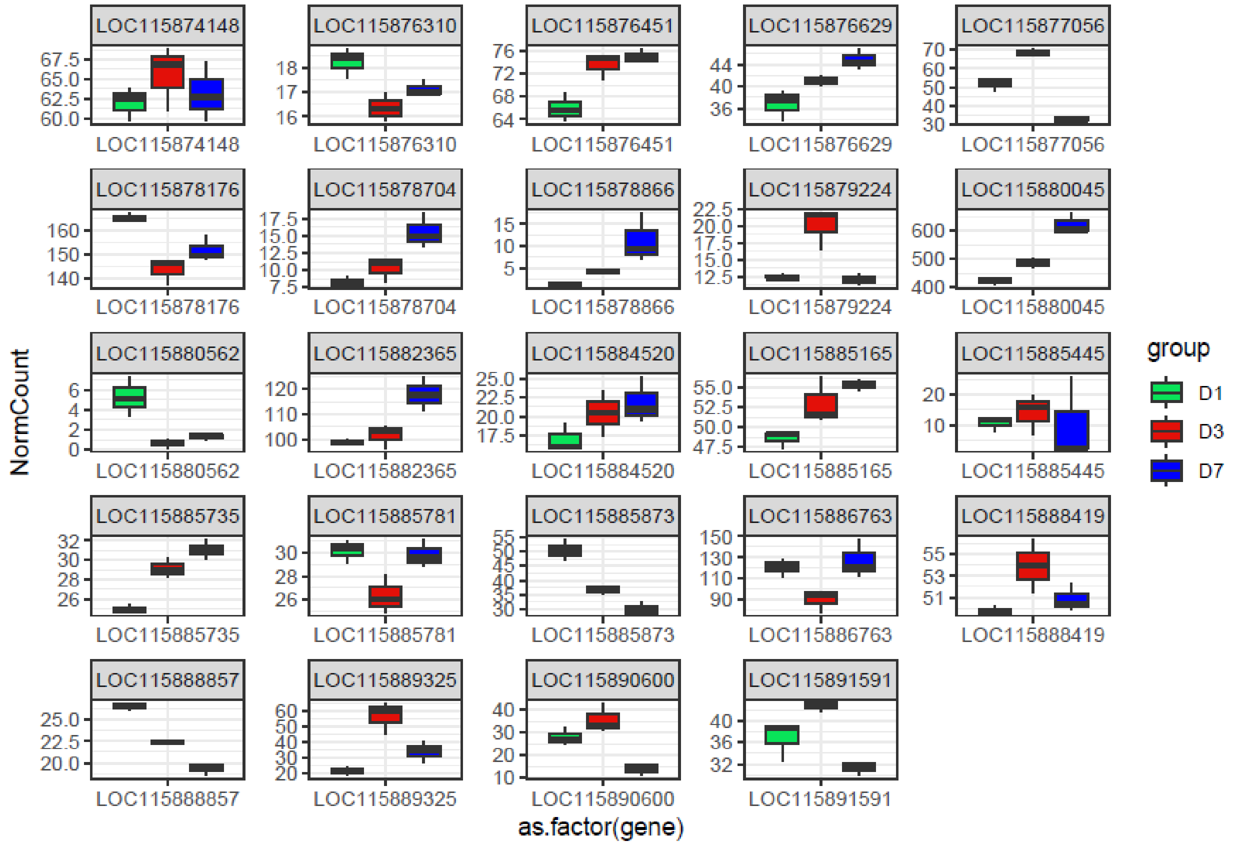


Figure S6D Normalized counts per million (CPM) of expressed IMD related genes in D1, D3 and D7 midgut.

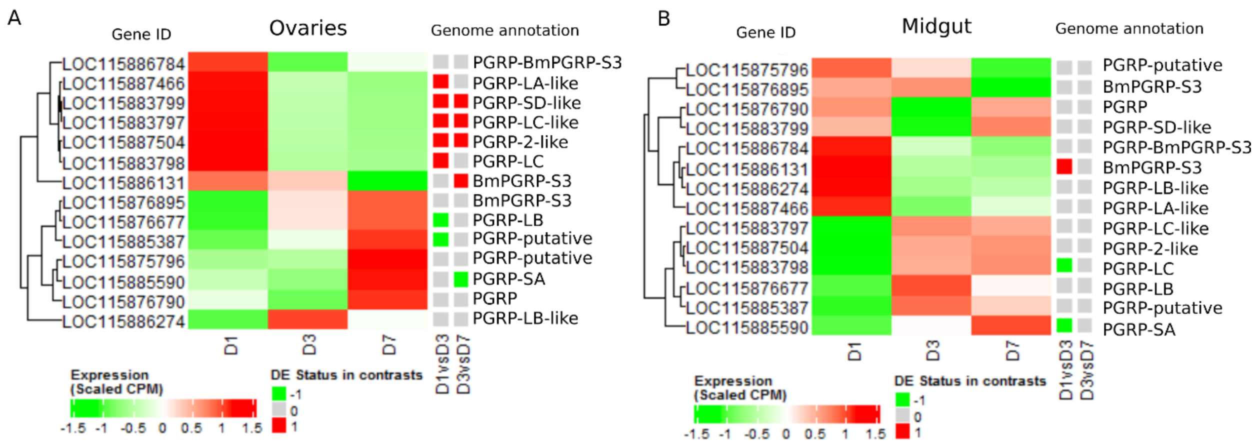


Figure S7. PGRP-gene-centered analysis in the germarium and midguts of D1, D3 and D7 adults. A-B. Heatmap of scaled CPM expression of genes in germarium (A) and midgut (B). Differentially expressed status between D1 and D3, and D3 and D7 is depicted with a coloured squared (red upregulation, green downregulation).

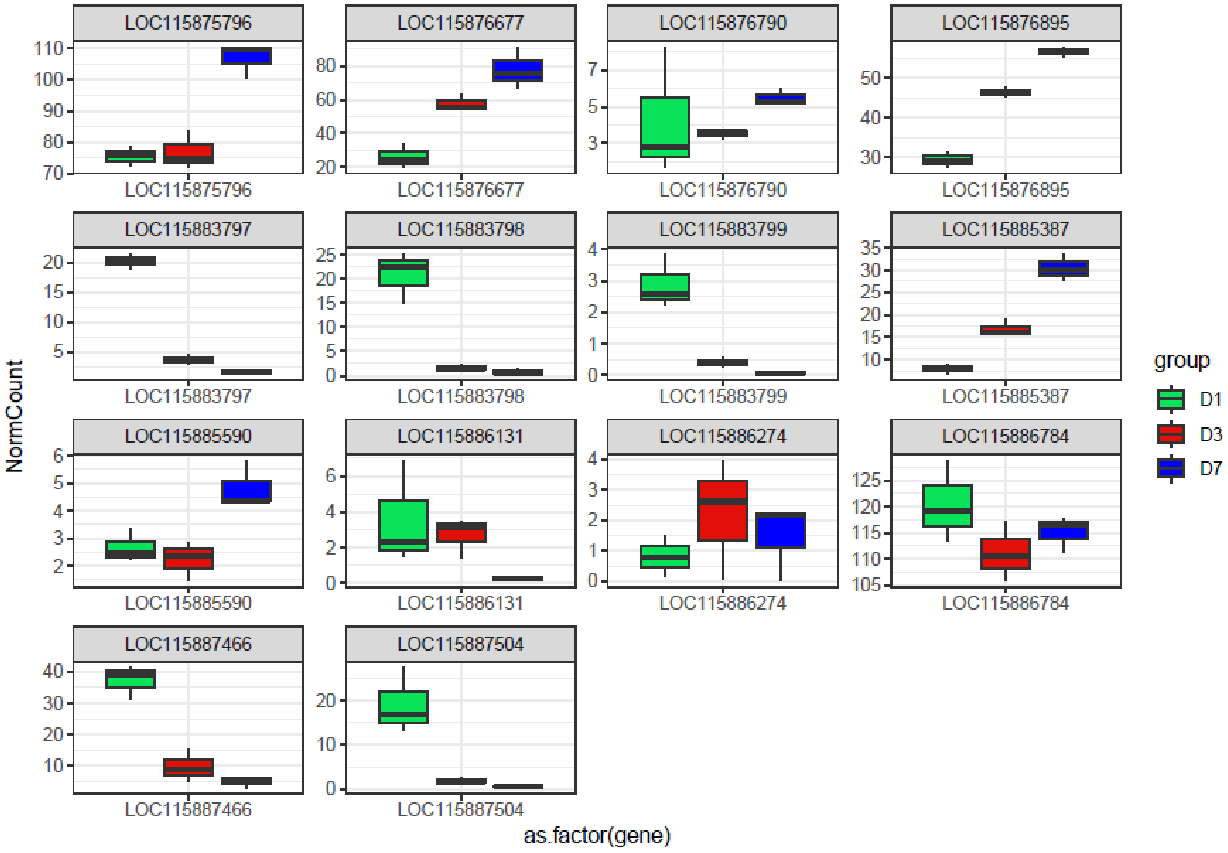


Figure S7C. Normalized counts per million (CPM) of expressed *pgrp* in D1, D3 and D7 germarium.

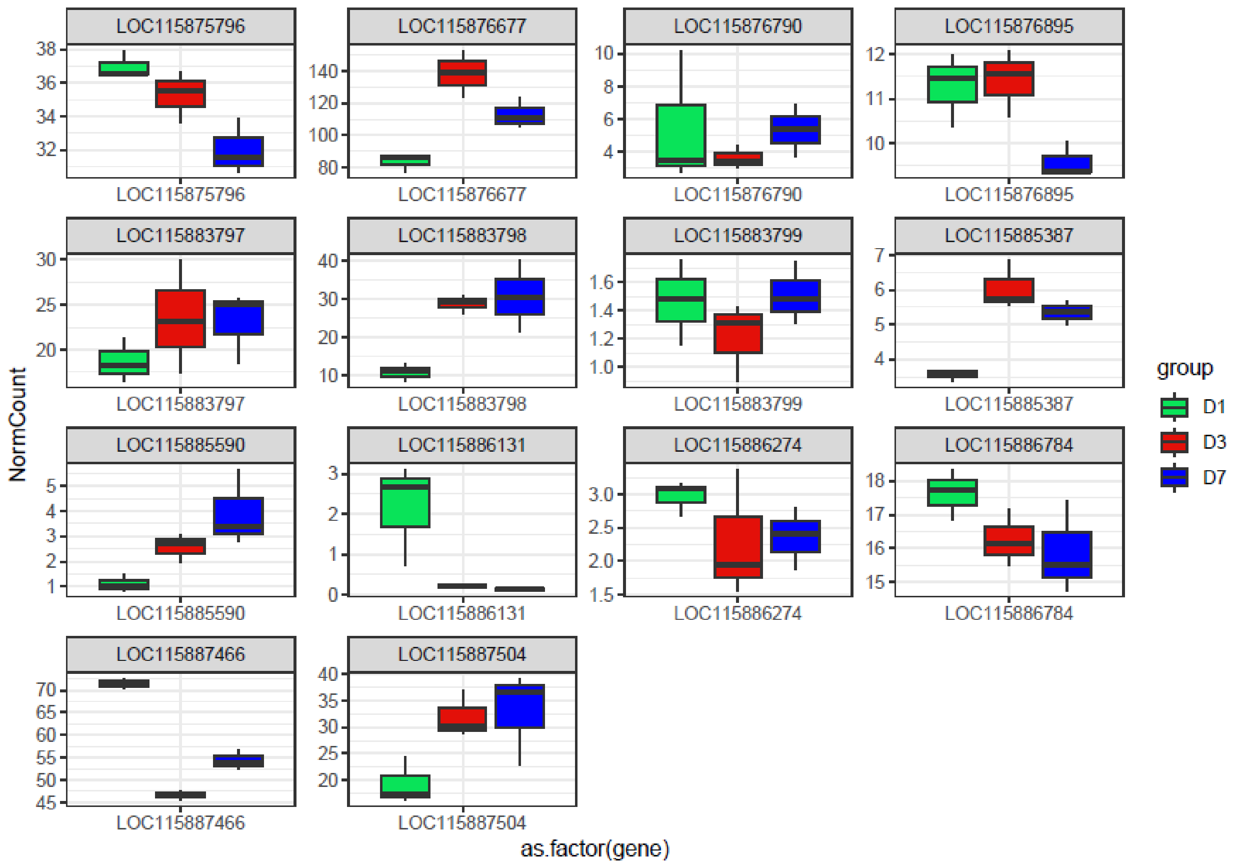


Figure S7D. Normalized counts per million (CPM) of expressed *pgrp* in D1, D3 and D7 midguts.
